# Generative Neural Spike Prediction from Upstream Neural Activity via Behavioral Reinforcement

**DOI:** 10.1101/2023.07.25.550495

**Authors:** Shenghui Wu, Xiang Zhang, Yifan Huang, Shuhang Chen, Xiang Shen, Jose Principe, Yiwen Wang

## Abstract

It is quite challenging to predict dynamic stimulation patterns on downstream cortical regions from upstream neural activities. Spike prediction models used in traditional methods are trained by downstream neural activity as the reference signal in a supervised manner. However, downstream activity is unavailable when neurological disorders exist. This study proposes a reinforcement learning-based point process framework to generatively predict spike trains through behavior-level rewards, solving the difficulty. The framework is evaluated to reconstruct the transregional spike communication during motor control through behavioral reinforcement. We show that our methods can generate spike trains beyond the collected neural recordings and achieve better behavioral performance.

---

Communication through neural spike signals is essential for complex brain function^1^. Neurological diseases or injuries that damage the transmission pathways can lead to dysfunction, such as memory impairments in Alzheimer’s disease or motor disabilities resulting from spinal cord injuries. A promising solution to restore functional loss is to construct an artificial information pathway to bypass the damage. For instance, one can build a biomimetic model that receives neural signals from upstream regions and predict neural activity in downstream regions. By delivering electrical stimulations to the downstream based on model predictions, such an approach can partially “rewire” the disconnected brain areas and induce similar functionalities^2, 3^. In fact, this has been applied in the memory prosthesis^4, 5^, which used point process models to predict spikes of the CA1 area as stimulation patterns from spike recordings of the CA3 area in the hippocampus. The prosthesis successfully improved the performance of human subjects in delayed image-matching tasks. For the neural pathway of motor control, researchers have interpreted movement intentions from the primary motor cortex and stimulated muscles^6, 7^ or spinal cord^8^ to restore motor functions of limbs for paralyzed individuals.

A key challenge for such an artificial information pathway is generating appropriate downstream neural activity. Prevailing spike prediction models^9-12^ were trained by supervised learning (SL) methods, which used recordings of downstream region spikes in pre-designed tasks as the ground truth of model outputs. This poses a challenge for patients with neural pathway damage, as they may have difficulties completing the task, and the recordings of downstream neural spikes may be either unavailable or abnormal^2^. Even if normal neural activity can be obtained, simply reproducing the recorded spikes might not be the optimal stimulation pattern for desired behavioral responses^13^. This is because the recorded neural spikes are typically obtained from a small cortex area over a limited time, thereby covering only a subset of the overall brain state. Meanwhile, SL-based models essentially estimate the distribution of recorded downstream activity conditioned on the upstream recordings, and thus cannot capture the downstream information beyond the recording regions that contributes to behavior. In other words, while the limited recordings provide a reference for supervised learning, they also constrain the models from exploring more effective solutions for behavioral rehabilitation.

An ideal model that functionally reconnects neural pathways should adapt its outputs without full knowledge of the target activity in downstream regions, similar to the neural plasticity of the brain when learning through trial and error. Therefore, we seek a generative spike prediction model that can surpass the statistics of recorded spike patterns in the target region. Reinforcement learning (RL) is a potential candidate for this solution, which explores all possible outputs for the best behavioral reward. RL algorithms have been applied to decode limb movements from neural spike counts in some brain-machine interfaces (BMIs)^14-17^. These studies predicted deterministic motor signals, such as position or movement direction. Thus, they are not suitable for predicting multi-channel neural spikes, which are high-dimensional and stochastic. RL models have also been implemented to predict point process events such as 911 crime incident calls and earthquakes^18-20^. However, these models only used self-history as inputs to predict the future, while the absence of a reliable history of neural spikes in the target region makes these methods inapplicable for spike prediction models.

Here we present the first RL-based point process (RLPP) framework for neural spike generation aimed to restore functional spike communication. The framework involves a spike prediction model that predicts downstream spike patterns from upstream spikes and learns from behavioral rewards. We evaluated the RLPP framework on the functional motor control pathway as a case study, using neural data from rats performing a two-lever discrimination task. With the proposed RLPP framework, the downstream spike trains generated by the spike prediction model exhibit similar patterns as the real recordings, leading to desired behavioral responses. These generated spike trains outperform those predicted by SL-trained models in the behavioral task, relying only on behavioral reinforcement and without needing ground truth. We further demonstrate that our approach can adapt to different settings of the behavior decoder with reasonable output spike patterns and robust behavioral performance. This relieves the dependence on a specific decoder or full knowledge of the neural pathway from the downstream region to the behavior. Besides, the RLPP-generated spikes have more distinguishable patterns than the real recordings under different movements. Detailed analysis suggests that our RL-based method utilizes more behavior-related information conveyed in the recorded upstream activity and generates spike trains beyond the recorded downstream spike structure.

## Results

### General architecture of RL-based spike generation

A reinforcement learning framework includes a model, an environment, and a learning algorithm. The model provides outputs to the environment in response to inputs, and the environment evaluates the model outputs and returns rewards. RL algorithms then update the model parameters to learn the input-output mapping that maximizes future rewards. The RL training is goal-directed and not constrained by any output ground truth. Thus, a spike prediction model trained under the RL framework can flexibly explore all possible patterns in the output space, provided the elicit rewards. The model output can be used as stimulation patterns for the downstream neural structure and further passed through the intact neural pathway to induce the subject’s behavior-level changes. The behavioral reward signal can then guide model learning (Fig. 1a). Note that the RL model does not intend to achieve the same downstream neural activity as the real recordings, which would be theoretically impossible given the fact that the downstream region also receives inputs from many other cortical areas. Instead, our RL-based method allows generating spike patterns beyond the real neural activities, as long as they can accomplish desirable behavioral tasks or restore desired functions.

**Fig. 1.**
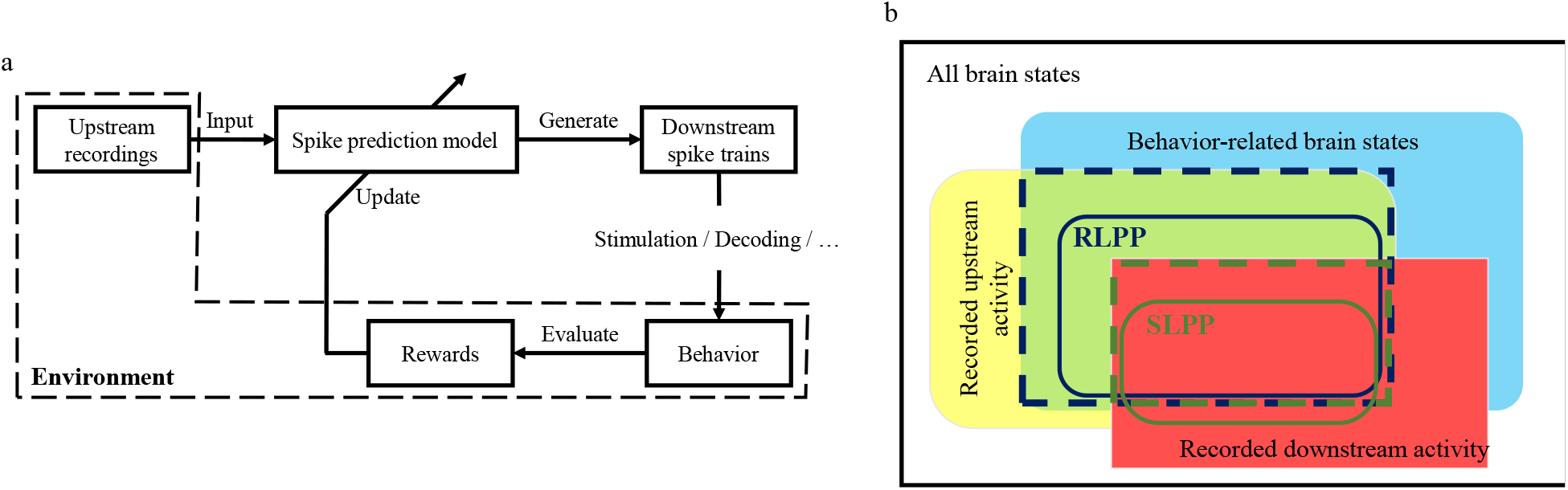
General structure and information flow of RL-based spike generation. **a**, General structure of reinforcement learning (RL)-based spike generation. A spike prediction model accepts input spike trains from upstream recordings and generate downstream spike trains as output. The output spikes can then lead to behavioral changes through electrical stimulations on the downstream region, decoding with a certain decoder, or other ways. By evaluating the behavior, we can update the spike prediction model with a reinforcement learning algorithm. The upstream regions and the pathway from generated spike trains to behavioral rewards can be viewed as an environment for the RL-based spike prediction model. **b**, A conceptual sketch for the information flow during the spike generation. Within the overall brain states, the yellow, red, and cyan blocks correspond to the brain states involved in recorded upstream activity, recorded downstream activity, and the behavior-related brain states, respectively. The supervised learning methods (SLPP, green box) can only capture the mutual information conveyed both in recorded upstream and downstream activities (dashed green box). The spike generation through behavioral reinforcement (RLPP, blue box) can explore within all task-related information encoded in upstream recordings (dashed blue box).

A conceptual diagram from the information perspective is given in Fig. 1b to illustrate the superiority of the proposed RLPP framework in behavioral performance. The neural activities recorded from the brain only reflect different subsets of the information conveyed in overall brain states, with some recordings being more related to the target behavior. In this way, supervised learning-based spike prediction methods, referred to as SLPP (green box), learn a restricted functional mapping from the recorded upstream (yellow block) and downstream (red block) neural activities conditioned by the time structure of the input. Obviously, the behavior-related information captured by the SLPP predictions would also obey the statistics of both recordings and the behavior (cyan block), shown as the dashed green box. In contrast, our RLPP method (blue box) explores all possible output patterns toward the behavior rewards, which is not conditioned by the input spike train structure. Mathematically, collected spike data are realizations of a random process, which has some statistical structure. Each realization has a different time structure that expresses the same statistical structure. Thus, the proposed architecture can utilize more abstract information (the statistics) from the upstream recordings for spike generation, not limited to the collected realization (dashed blue box). In other words, we expect the RLPP will outperform the SLPP in behavioral tasks. While the SLPP models are upper-bounded by the model generalization obtained from upstream and downstream random process realizations, the upper bound of the RLPP model would only be determined by the behavior-related information (statistics) encoded in the upstream recordings. In essence, our study implements an online, multi-channel, spike-to-spike prediction using only behavioral feedback, delivering a model that can adaptively generate *spike ensembles* and participate in neural communication as a “silico cortical circuit”.

### Data preparation and framework implementation

We show the capability of the RLPP framework on spike data recorded from the medial prefrontal cortex (mPFC) and the primary motor cortex (M1) of six male Sprague Dawley (SD) rats. These two regions play essential roles in decision making and motor control, with the mPFC being the upstream region and the M1 being the downstream^16, 21-23^. Each rat was surgically implanted with two electrode arrays in the left hemisphere, one in the M1 and the other in the mPFC. The rats were placed in the behavior box and free to move while performing a two-lever discrimination task^24^. Neural spiking activities were recorded while the rats made their choices and pressed one of the two levers in response to a random sequence of audio cues in two different pitches (Fig. 2a). The start cue for the coming trial would be given 4-7 seconds after the previous trial ended. We collected about 200 successful trials over a 40-minute session for each rat and sorted the recorded neural signals offline. All neural and behavioral data were binned with a time resolution of 10 ms. M1 neurons were selected by finding a minimum set to reconstruct the rats’ behavior (see Methods). For each rat, spike trains of 13-25 mPFC neurons were recorded as model input, and the spike trains of 2-10 M1 neurons were selected for model prediction.

**Fig. 2.**
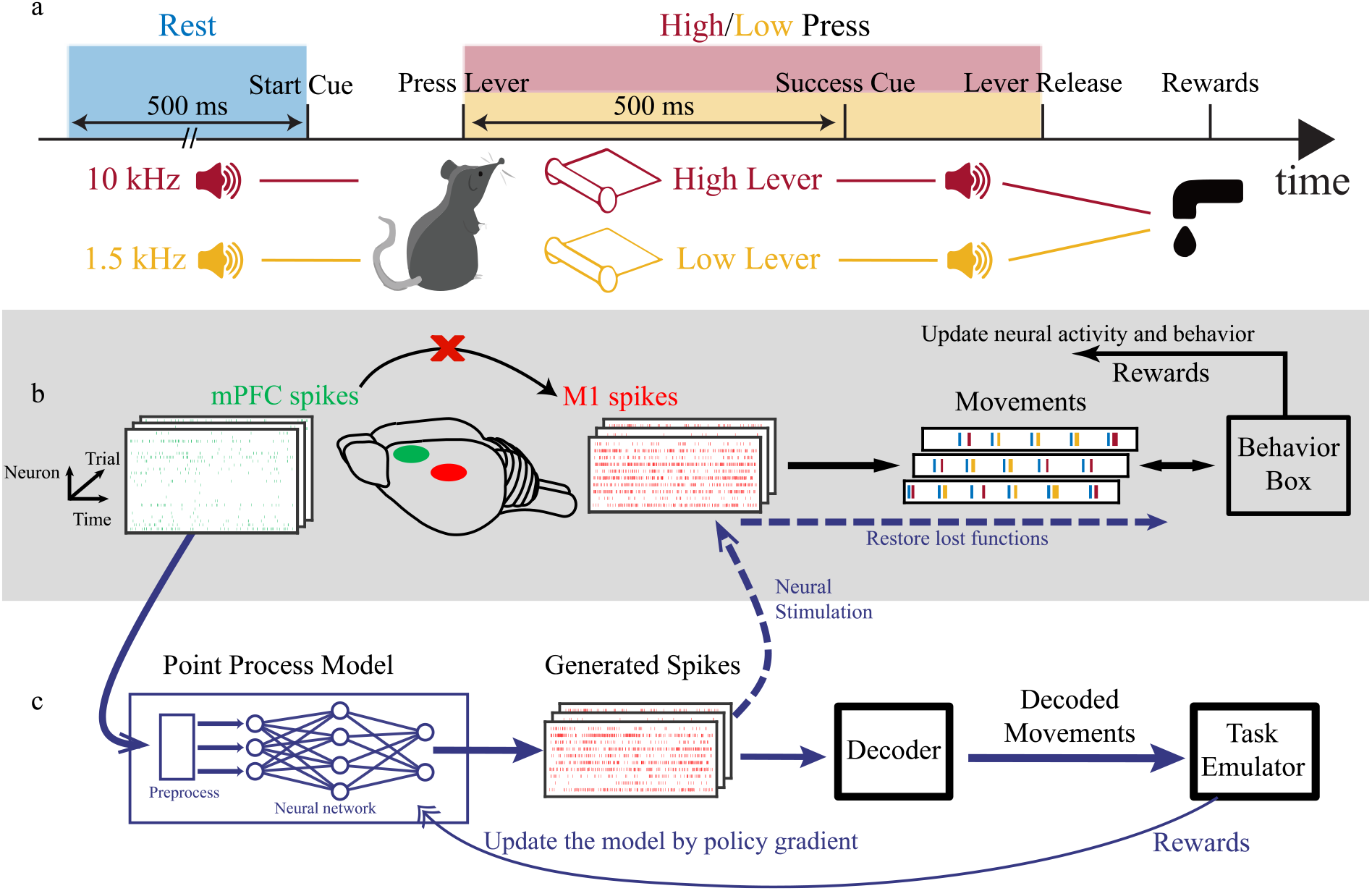
Experimental paradigm and framework implementation. **a**, Paradigm of the two-lever discrimination task. Rats were placed in the behavior box while neural activities were recorded from the primary motor cortex (M1) and the medial prefrontal cortex (mPFC) using implanted electrode arrays. Rats could move freely in the box and were required to press and hold the corresponding lever according to a sequence of randomly generated audio cues. We define the 500-ms duration before the start cue as the rest state of rats, and the duration between lever pressing and releasing as the high/low press states. These states are used to label the rats’ movements during the task. **b**, The information pathway of motor control and learning (black arrows). The mPFC makes decisions based on the environment and passes information to the M1, which executes movement signals through the neuro-muscular system. Rats learn to adjust their movements through the reward feedback during interactions with the behavior box. A disconnection from the mPFC to the M1 may lead to abnormal M1 activity and failure in the task. **c**, Implementing the reinforcement learning point process (RLPP) framework (blue arrows). A point process spike prediction model generates the M1 spikes from mPFC spikes. The generated M1 spikes are decoded into movements through a decoder and evaluated by the task emulator. Then, the model is updated according to the rewards by the policy gradient algorithm without using real recordings of M1 spikes. Model prediction-based neural stimulations on the M1 neurons can bypass the damaged neural pathway and restore the motor function through the intact neuro-muscular system (blue dashed arrows).

We implemented the proposed RLPP framework to rewire the functional pathway of neural spikes from mPFC to M1 as a case study. When the connections between mPFC and M1 were not fully functional, the normal M1 neural activity would be affected, leading to a failure in the behavioral task (Fig. 2b). To address this issue, a point process spike prediction model learns a functional mapping from the neural spikes of the recorded mPFC neurons to the firing probabilities of the selected M1 neurons, and generates M1 spikes through sampling from the firing probabilities by inhomogeneous Bernoulli processes (Methods and Fig. 2c).

The model is trained using behavior-level responses in a RL framework. We first decode the generated M1 spikes into movements to mimic the rats’ responses induced by the stimulations on M1 neurons. Rewards are given to the spike prediction model when the movements can accomplish the task. We implemented a policy gradient algorithm for the multi-neuron point process to optimize the spike prediction model. The movement decoder is trained before the spike prediction model, using the recorded M1 spikes as decoder inputs and the rats’ behavior as output targets. Although this study did not involve actual subject stimulation, utilizing the decoder as a proxy instead, we believe the decoder is still practical and helpful as a controlled environment for real stimulation. We elaborate on this aspect in the Discussion section. In addition, we examined the generalization ability of the RLPP framework by using first a manual-designed decoder and later a more realistic cross-subject decoder.

For comparison, we also trained spike prediction models through supervised learning-based point process (SLPP) methods^9, 11^. These models have the same structure and input as those used in the RLPP framework, optimized by minimizing the point process log-likelihood between model predictions and the recorded M1 spike trains. We trained the RLPP and SLPP models under five-fold cross-validation on each rat’s ata and characterized the recorded and predicted spikes using the peri-event firing probability and mutual information. The predicted M1 spike patterns were further used as input to the movement decoder to perform the two-level discrimination task. The behavioral level performance of the trained models was measured by the time-bin success rate and trial success rate.

### Can RLPP generate behavioral-relevant spike trains?

To answer this question, we compared the raster plots and peri-event firing probabilities of predicted spikes to those of real M1 spike recordings. Fig. **3**a-c shows the modulation of six M1 neurons, which are part of the input in behavior decoding.

**Fig. 3.**
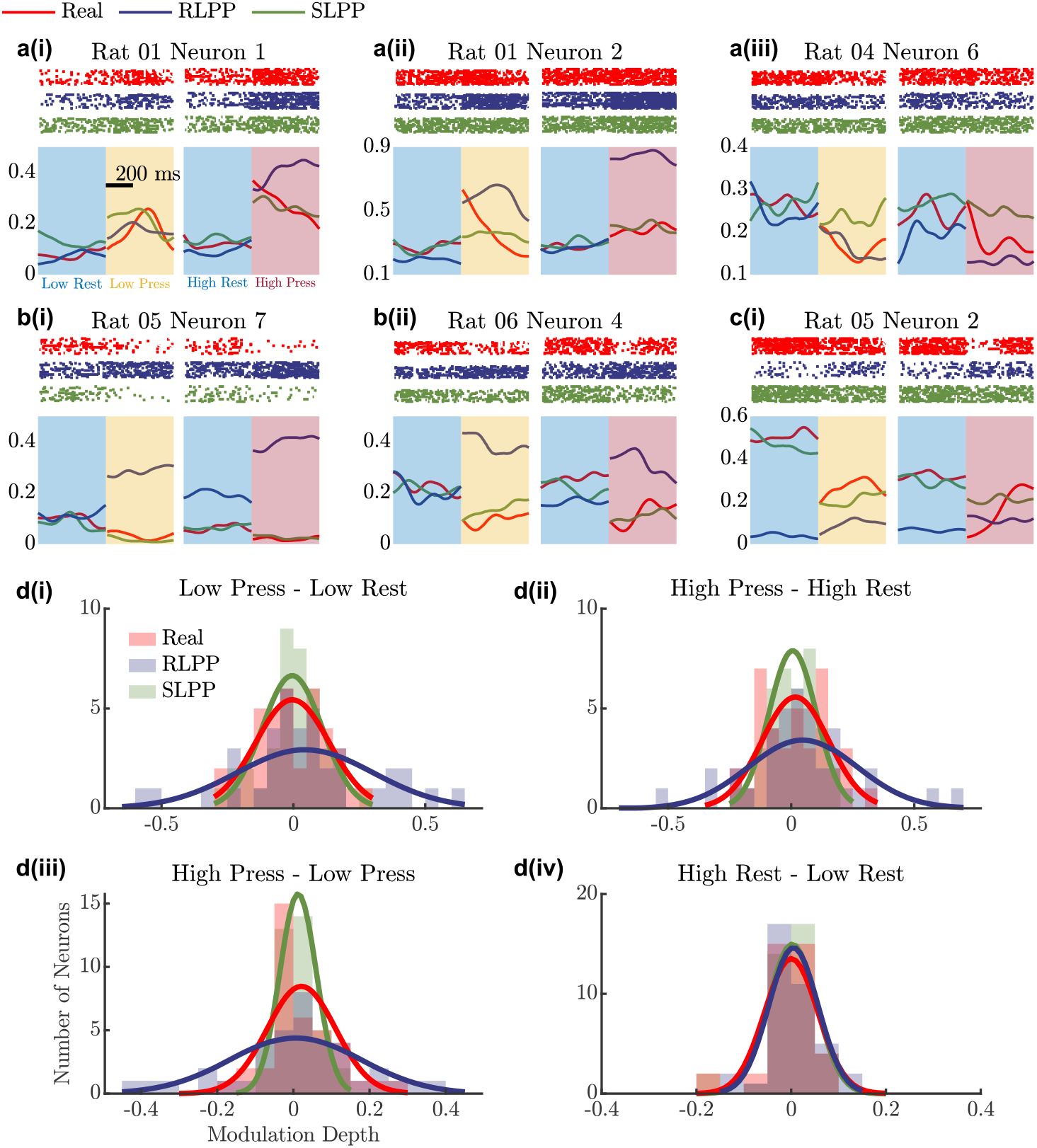
Neural modulations of M1 spikes predicted by different methods with respect to movement states. **a-c**, Modulations of individual neurons. For each subplot, the left half (cyan and yellow blocks) compares the neural activity during the rest state (500 ms before the start cue) and low press state (500 ms during the lever pressing) of the low lever trials. The right half (cyan and pink blocks) compares the neural activity during the rest state and high press state of high trials. The raster plots show the spike trains across multiple trials. The curves show the peri-event firing probability. The horizontal axis is the time index. The vertical axis for raster plots is the trial index, and for the curves is the firing probability normalized across trials. The red, blue, and green colors represent the M1 activity of real recordings, RLPP predictions, and SLPP predictions, respectively. **a** shows three cases that RL-predicted M1 spikes can track and enhance the modulations; **b** shows two cases that RL-predicted M1 spikes can generate spike patterns with new modulations not observed in real data; and **c** shows a rare case that RL-predict spikes do not modulate with the movements. **d**, The histogram and estimated distribution for modulation depth of real and predicted spikes across M1 neurons from six rats. The modulation depth evaluates the changes in average firing probabilities of one M1 neuron during two different movement states. The distribution of the modulation depth is estimated by Gaussian functions. The horizontal axis is the modulation depth. The vertical axis is the number of neurons. The red, blue, and green colors represent the modulation depth of real recordings, RLPP predictions, and SLPP predictions, respectively. The title of each subplot shows the two movement states compared in the subplot. Overall, the RLPP predictions have higher modulation depth, while the SLPP predictions have lower modulation depth when compared with real M1 spikes.

In general, the RLPP model successfully generated M1 spike trains with task-related modulations. In Fig. 3a, the RLPP predictions show different firing probabilities among the rest, low press, and high press actions. These modulations in RL-predicted spike trains are similar to those in real spike trains during task execution, indicating that the RLPP model effectively tracks and even enhances the discrimination in real spike trains. Moreover, the RL predictions are within or partially within the mean ±1 standard deviation of the real spike trains across trials (not shown), despite not using any M1 recordings. In contrast, SL-predicted spikes show little difference in Neuron 2 of Rat 01 when pressing the low lever and Neuron 06 of Rat 04 when pressing the high lever (Fig. 3a(ii) and (iii), respectively).

Fig. **3**b gives two examples of RLPP models finding modulations not observed in the collected data. Although the recorded spikes of both neurons are negatively modulated with lever pressing, the RLPP predictions show strong positive modulations on these two neurons. Fig. **3**c depicts one of the rare cases where the RLPP predictions may not be modulated with movement information on the given neuron, while the real spike data does. Although the RL-predicted spikes for these neurons have different modulations compared to the real recordings, they do preserve task success.

We also analyzed the modulation depth of neurons, defined as the difference between average firing probabilities during two different states. Fig. **3**d shows the histograms and distributions of the modulation depth for real recordings, RLPP, and SLPP models. For example, the real recordings exhibit few differences in the average firing probability during the rest state before low or high trials, showing a small range of modulation depth (the red curve and bars in Fig. 3d(iv)). This is as expected since the rest state should be similar regardless of the following start cues. RLPP and SLPP models capture this property, as the blue (RLPP) and green (SLPP) bars/curves are almost identical to the red ones in Fig. 3d(iv).

Real recordings for other movements show a large modulation depth range (Fig. 3d(i-iii)). The firing probability of some neurons significantly change according to the movements. In comparison, RLPP-predicted spikes have a larger range of modulation depth than real spikes, while the SLPP predictions have a smaller range. Such a larger modulation depth indicates better separability in RL-predicted spike patterns, which leads to better movement decoding and higher behavioral performances, as shown in the next section. The differences between RLPP and SLPP predictions will be quantified with an information theoretical analysis, as shown in the last section of Results.

### Can RLPP-generated spikes lead to realistic behavior?

Here, we would like to present the generated spike trains in the time domain. Fig. **4** shows the spikes and firing probabilities of two M1 neurons (Fig. 4a) alongside the movements (Fig. 4b) within 6-7 trials. The rhythmic firing patterns of both neurons in the real data suggest that these two neurons are positively modulated with the press movements. The predictions of the RLPP and SLPP successfully follow the trend, such as the increase in Neuron 1’s firing probability in 2-3 seconds (Fig. 4a(ii)). We can also see that Neuron 2’s RL-predicted firing probability is significantly larger during pressing than the real neuron firing, while the SLPP predictions show little fluctuations.

**Fig. 4.**
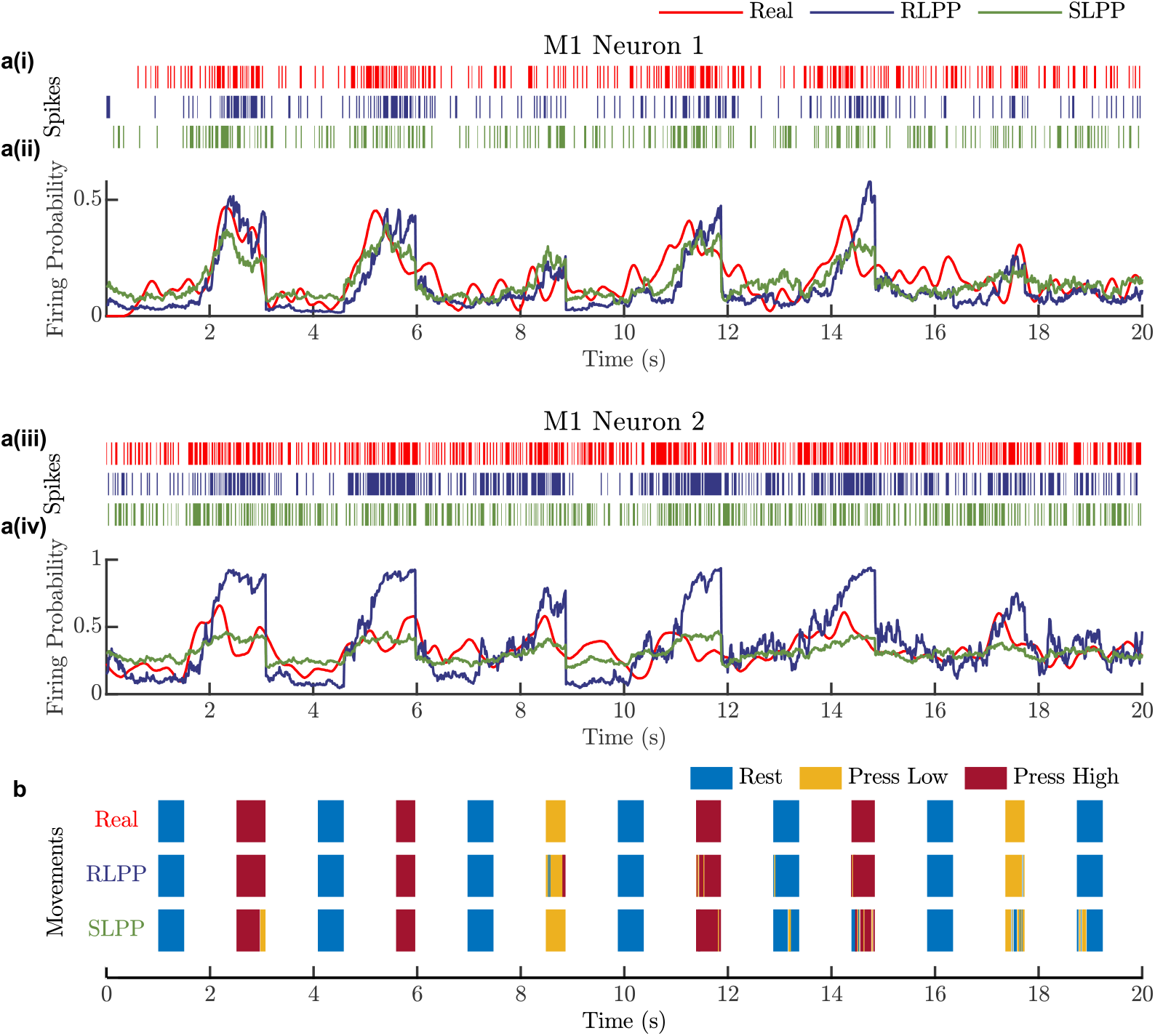
Predicted M1 spikes and decoded movements of Rat 01. **a**, Spike trains (i and iii) and firing probabilities (ii and iv) of the two selected M1 neurons. The red spike train is the real recordings of the rat performing the behavior task, and the firing probabilities (red curve) are smoothed from the recorded spikes by a Gaussian kernel. The blue and green curves represent the predicted firing probabilities from the models trained by the RLPP and SLPP, respectively, with the corresponding predicted M1 spike trains depicted in the blue and green spikes trains. **b**, Real movements (first row) of Rat 01 during the task and decoded movements from RLPP (second row) and SLPP (third row) predicted M1 spikes. The blue, yellow, and red bars represent the rest, press low, and press high movement during the task, respectively. All plots share the same horizontal axis as the time index. The vertical axis of a(ii) and a(iv) is the firing probability. Data shown in the plots are a segment of the test set of Rat 01.

These observations concur with Fig. 3a(ii), where the RLPP-predicted M1 spikes have a larger modulation depth. We also compared the decoding performance of the predicted spikes in the lever discrimination task. The first row of Fig. 4b shows the real movements performed by the subject during the task, serving as the desired decoding targets. The other two rows show the movements decoded from the predicted spikes, which have similar patterns to the targets in the first row. Moreover, the model predictions in certain time bins may be translated into wrong movements and violate the task requirements, shown as the color strips. For example, the SLPP predictions failed to generate high press movement constantly during 14-15 seconds. The RLPP predictions made fewer mistakes than the SLPP predictions in the task, indicating that the proposed RLPP framework can lead to better behavior-level performance.

### Statistical behavior-level evaluation of generated spikes

The utility of our RL-based spike prediction framework was further examined by quantifying the statistical performance of the decoded movements. The time-bin success rate, defined as the ratio of time bins when the decoded movements matched the requirements of the behavioral task, was used to evaluate the average performance across time. Additionally, the models were compared on a trial basis, where a trial was considered successful when the correctly decoded actions dominated the time instances within the trial (over 70% of the time bins). The trial success rate was defined as the ratio of successful trials over all trials in the test set. The results in Fig. 5 demonstrate that the RLPP predictions significantly outperformed the SLPP predictions on both metrics. This may be because the RLPP framework is designed to accomplish the task and is not constrained by the recorded M1 spikes, leading to better performance in motor control. Overall, these results demonstrate that the RLPP model can effectively generate task-related M1 spikes that can be decoded into realistic movements.

**Fig. 5.**
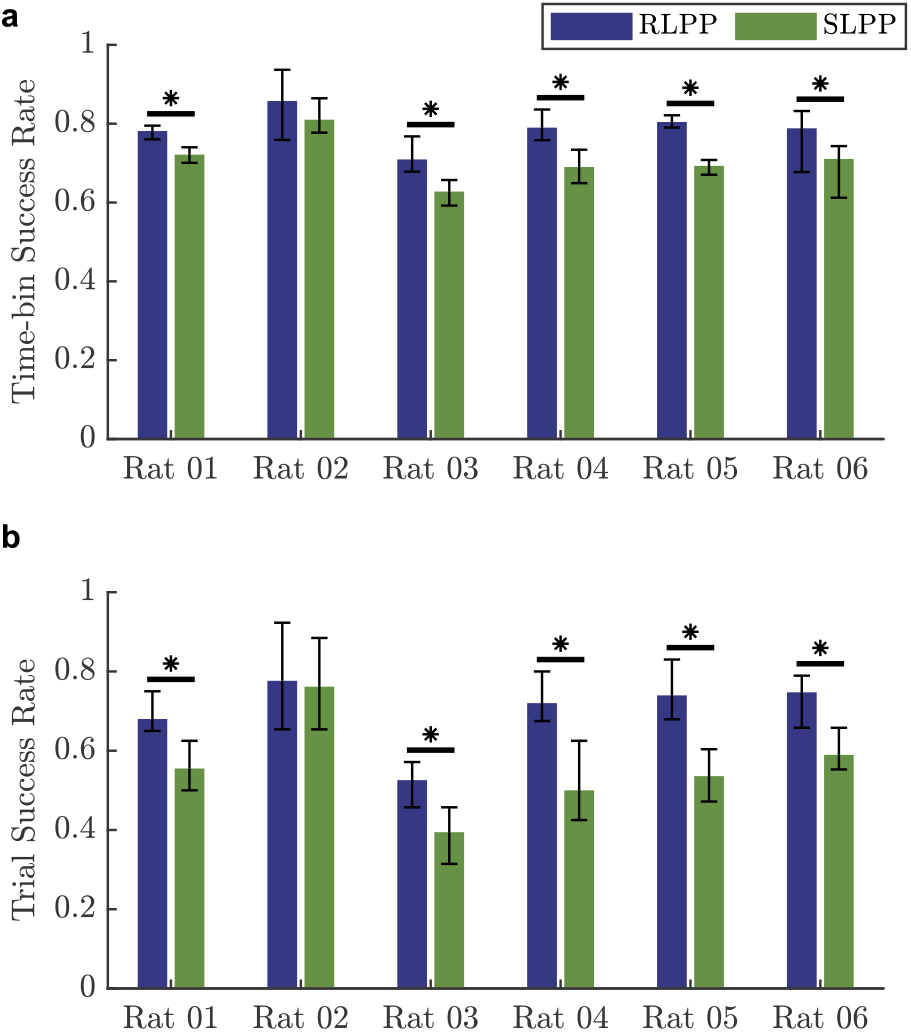
Comparing the behavior-level performance of RLPP and SLPP. **a** and **b** show the time-bin success rate and the trial success rate of the movements decoded from the predicted M1 spikes, respectively. Each bar shows the mean and range of the performance across the test sets of five-folds for each rat. The stars above each two bars mean ∈ ∈ 0.05 by the right-sided Wilcoxon signed rank test for paired samples. Our results show that RLPP outperforms SLPP for all six subjects on both metrics, suggesting that the RLPP framework can lead to better behavioral performance.

### Extending RLPP to manually designed or cross-subject behavioral decoders

Here we show that the RLPP framework is not restricted to a specific decoder trained for each subject, instead, can predict behavior-related downstreram spikes with different decoder settings and still complete the original behavioral task. We selected Rat 02 and Rat 05 as two examples to show the capability of RLPP. In the first part of this section, we manually designed a decoder, which assigns different preferred directions to “artificial” M1 euro s. In this two-lever discrimination task, we generated four “artificial” euro s to cover different potentials, where Neuron 1 (2) is positively (negatively) modulated with low press, and Neuron 3 (4) is positively (negatively) modulated with high press. The results are illustrated in Fig. 6a-h, which reveal that the RLPP model successfully shapes the output spike trains to fit the decoder settings and accomplish the lever-discrimination task. For example, predictions on Neuron 1 have higher firing probabilities than Neuron 2 during 15-16 seconds in Fig. 6b, resulting in low press movements, and Neuron 3 is more active than Neuron 4 during 12-13 seconds for high press movements in Fig. 6c. Besides, Neuron 1 and 3 tend to have low firing probability during the rest movements, while Neuron 2 and 4 will have relatively higher firing. An interesting finding is that some “artificial” euro s have similar atter s ith the real M1 recordings of selected neurons, shown in Fig. 6d and h, although no M1 information is used in decoder design and RLPP training.

**Fig. 6.**
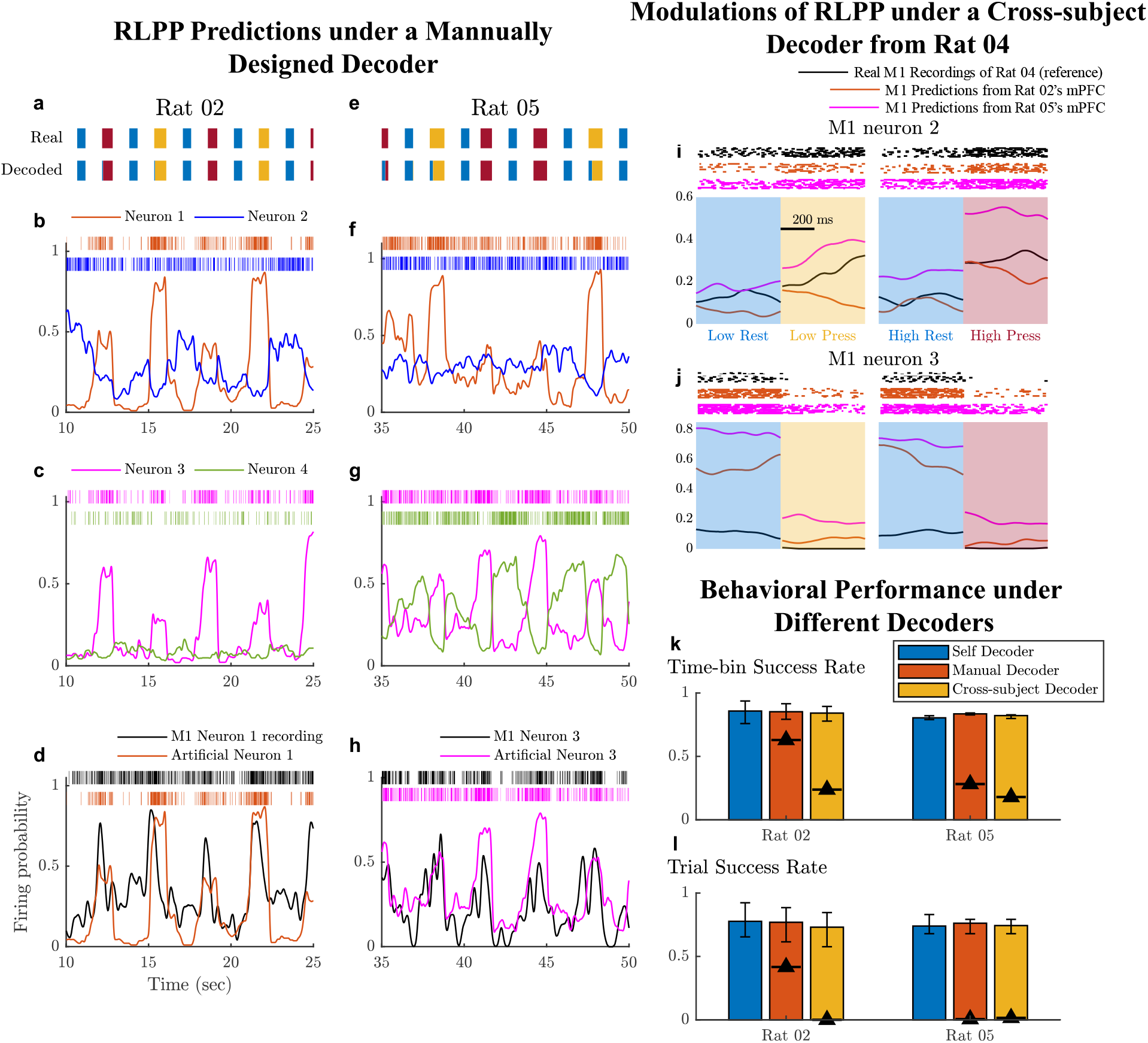
RLPP models perform well under different decoder settings. **a-h** illustrate the prediction and decoding results under a manual-designed decoder, where **a**-**d** correspond to Rat 02 and **e**-**h** correspond to Rat 05. The x-axis for **a**-**h** are the time index, and the y-axis for **b**-**d** and **f**-**h** are the firing probability. **· a** and **e**, Comparison between the real movements of the rats during the task and the movements decoded from the RLPP model with the designed decoder. The decoded movements are similar to the real movements with few errors. **· b**-**c** (**f**-**g**), RLPP-re icte firi robabilities a s i es of the four “artificial” euro s, sho b four colors. The curves sho the firing probabilities, and the bars show the neural spikes. The predicted neural activities show rhythmic patterns and clear modulations as the decoder designed. **· d** and **h**, Com ariso bet ee the recor e activities of o e real M1 euro a the re ictio s of o e “artificial” euro. The black color corresponds to the real recordings of M1 neurons, and the other color represent the RLPP predictions under the designed decoder. The model predictions show similar patterns as the real recordings of some selected M1 neurons, even no M1 recordings are used during the entire training process. **i**-**j** illustrate the modulation of RLPP predictions from mPFC activities of Rat 02 and Rat 05 under the decoder from Rat 04. The layout is the same as the subplots in Fig. 3. The black spikes and curves show the raster and modulation from the real recordings of Rat 04, and the orange and pink color correspond to the RLPP predictions. The M1 neuron 2 (3) is positively (negatively) modulated with the press movements. These features are captured by the RLPP models under the cross-subject scenario. **k**-**l** illustrate the behavioral level statistical results of RLPP model under different decoder settings with same mPFC inputs. The blue, orange, and the yellow colors correspond to the performance of RLPP with the self-decoder, the manual-designed decoder, and the cross-subject decoder of Rat 04, respectively. Each bar shows the mean value across the test sets of five folds, and the error bars show the range of the performance. Thus, the blue bars have the same values as the corresponding bars in Fig. 5. The black triangles label the average performance by the real M1 recordings of Rat 02 or Rat 05 with different decoders. The performance of RLPP is robust to the decoder settings, and greatly improved the behavioral performance of real recordings.

In the second part, we created a cross-subject learning scenario, where the spike prediction model accepts the mPFC activities of Rat 02 during the behavioral task, but the predicted M1 spikes are translated by the decoder of Rat 04. Rewards are given to the prediction model when the decoded movements accomplish the task that Rat 02 is performing. We also tested the same setup on Rat 05 with at 04’s decoder. The decoder of Rat 04 has seven M1 neurons as inputs. Fig. **6**i-j show the modulations of the predictions on two of the seven M1 neurons. The real recordings of Neuron 2 (3) are positively (negatively) modulated with the press movements, while the RLPP models also generate spike patterns with the same modulations from the mPFC recordings of both rats.

A comparison of statistical behavior-level performance is given in Fig. 6k-l. The task can barely be accomplished with the inputs of the real recordings of M1 neurons fed to unmatched decoders (shown by black triangles), whereas the performance of RLPP is robust to decoder settings, and significantly outperforms the real recordings. These results suggest that the proposed framework can be extended to various decoder settings and application scenarios.

### RLPP-generated spikes are more related to the desired behavior

The separability of M1 spike trains can be illustrated through the t-distributed stochastic neighbor embedding (t-SNE) technique^25^, a nonlinear dimensionality reduction method widely used to visualize high-dimensional data. We used the M1 spike train history at each time bin (input of the movement decoder) as data samples and applied t-SNE to project the data into a 2-D space. The data samples were labeled according to the real movements at the corresponding time bin.

The results of three rats are shown in Fig. 7. The real M1 recordings of Rat 02 and Rat 04 are clustered into three groups, with each cluster corresponding to a specific movement. No clear clusters are observed from the M1 recordings of Rat 03. However, SLPP predictions on Rat 04 show few differences between low and high press movements, suggesting that the SLPP model may miss some information in M1 activity when learning from the ground truth. In contrast, the RLPP-predicted M1 spikes can be clustered into three movement-related groups in all three rats. This indicates that the RLPP framework can capture the fine detail of predicted spikes for discriminability, which can improve the behavior-level decoding performance.

**Fig. 7.**
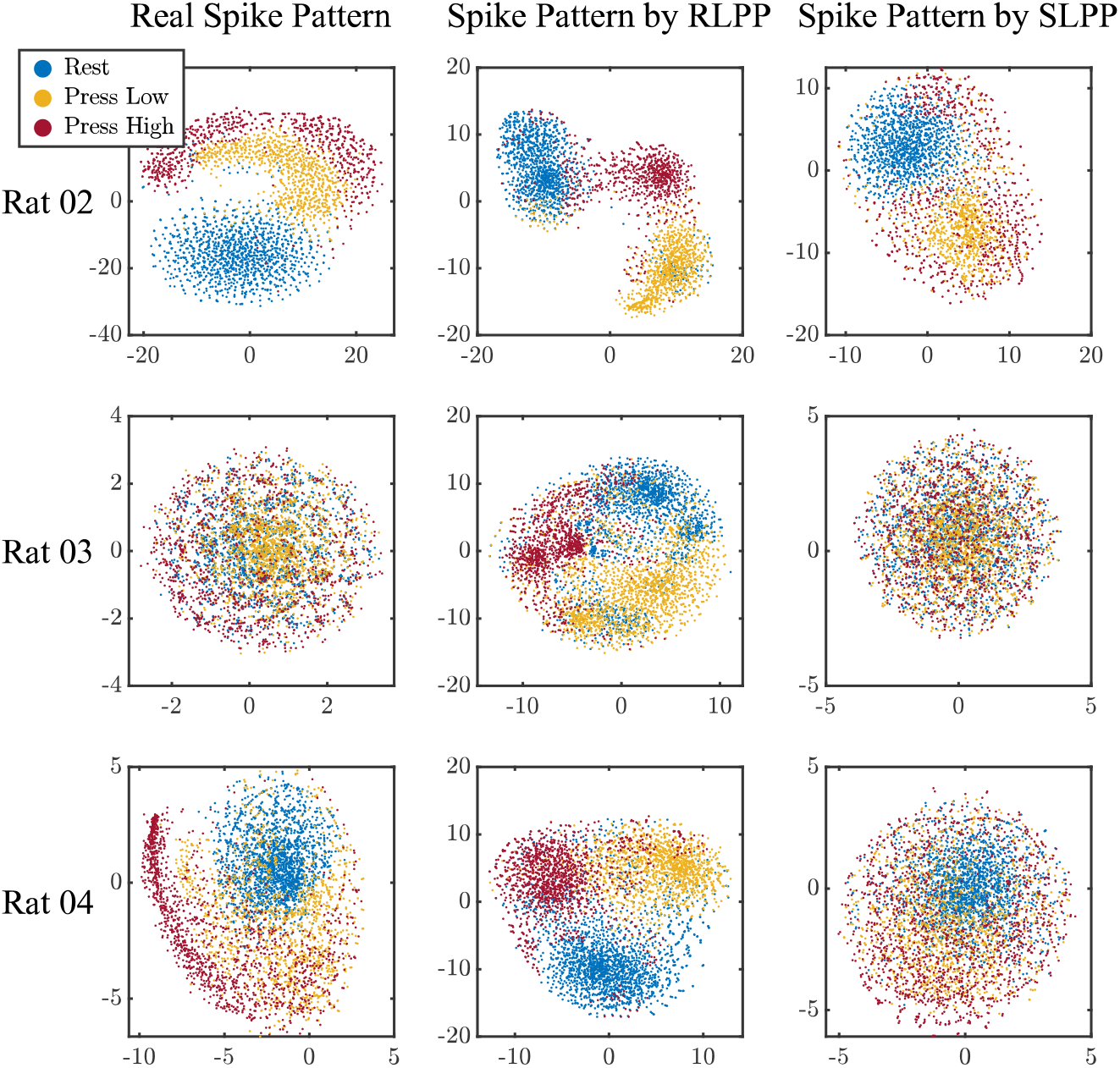
Visualizing M1 spike ensembles using t-SNE. The figure presents results from three rats, with each row corresponding to one rat. The M1 spike ensembles are projected into a 2-D space by t-SNE for visualization. The three columns display the results of real spikes, RLPP predicted spikes, and SLPP predicted spikes. The color of each data point represents the movements of the rats in real data at the corresponding time bin. Compared with the real spikes and SLPP predictions, the RLPP predictions exhibit better distinguishability on all three rats.

The relationship between model-predicted spike trains and behavior can be quantified by mutual information (MI), which measures the information shared between two stochastic processes. Here, we show that the RLPP model utilizes more movement-related information than SLPP through the MI values of the real M1 spikes, the SL-predicted spikes, or the RL-predicted spikes with respective to the real movements for each M1 neuron. Results are illustrated in Fig. 8a, where MI values for the same neuron are connected by lines. Most lines between the real and SLPP groups go from top-left to bottom-middle, as the SL-predicted spikes have significantly smaller MI values than the real M1 spikes. This suggests that the SLPP model did not fully capture the motor-related information encoded in the recordings of selected M1 neurons. The RLPP predictions have significantly larger MI values than the SLPP predictions, which means the RLPP-generated spike trains encoded more behavior-related information. No significant differences are found between the RLPP and real recording groups. However, we still see many neurons with higher MI values in RLPP predictions (labeled by triangles) than in real recordings, in line with the neurons shown in Fig. 3b, which demonstrate a larger modulation depth in the RLPP predictions.

**Fig. 8.**
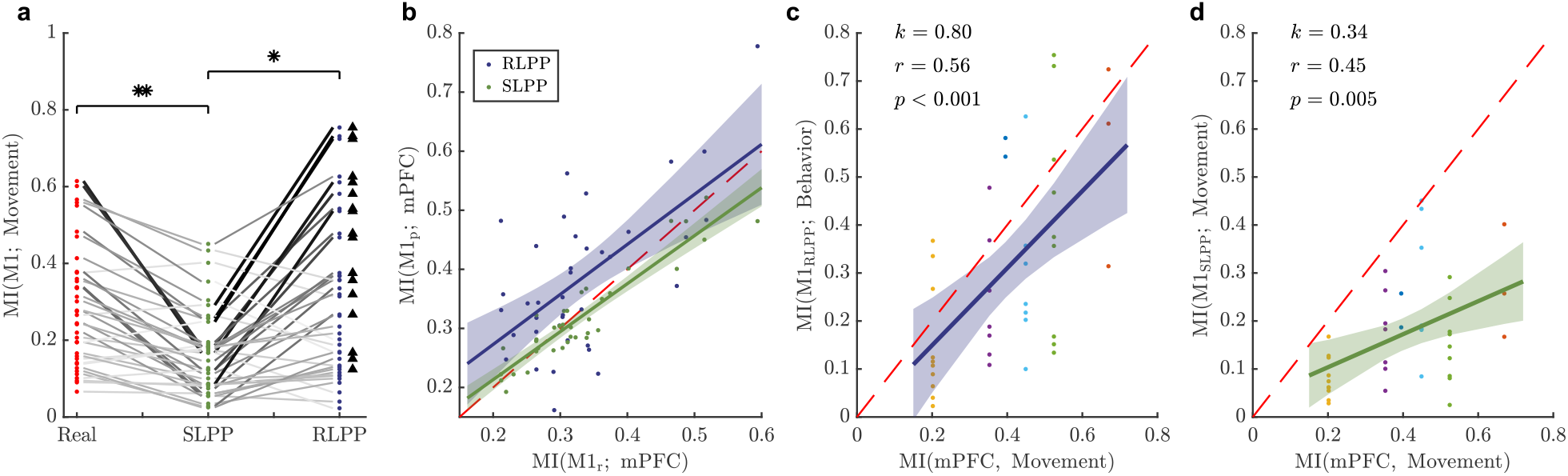
Information analysis for spike prediction models. **a**, Mutual information (MI) between real movements and M1 spikes from real recordings (red), RLPP predictions (blue), or SLPP predictions (green). The x-axis labels the source of M1 spikes, and the y-axis is the MI values. Each data point represents one M1 neuron, where the values for the same M1 neuron are connected by black lines. SLPP predicted M1 spikes have significantly smaller MI values with movements compared with real M1 spikes (*p* < 0.001) and RLPP predictions (*p* < 0.001) by the one-sided paired t-test. Predictions on some M1 neurons by RLPP show higher MI values (labeled by triangles) than real recordings, while no significance is shown by the one-sided paired t-test. **b**, MI values between the spike trains of two cortex regions. The x-axis is the MI value between real recordings of the selected M1 neurons (M1_*r*_) and the recorded mPFC neurons. The y-axis is the MI value between predicted M1 spike trains (M1_*p*_) and the recorded mPFC neurons. Each data point represents one M1 neuron, and the blue and green colors correspond to the RLPP and SLPP predictions, respectively. The blue/green line shows the results of linear regression on RLPP/SLPP predictions, and the shaded area indicates the 95% confidence intervals. For SLPP predictions, the correlation coefficient *r* = 0.∈, and the *p*-value of the linear regression is less than 0.001, indicating a strong and significant linear relationship between the compared MI values. The dashed diagonal line in red corresponds to the situations when the predicted and recorded spike trains of the same neuron have the same MI values with mPFC spikes. The MI values between SL-predicted spikes and mPFC spikes are significantly smaller than the MI values between real M1 spikes and mPFC spikes (*p* < 0.05, one-sided paired t-test), and 61% of the neurons have smaller MI values, corresponding to the green points below the red line. For RLPP, the MI values are significantly larger (*p* < 0.005), and 74% of the neurons have larger MI values (blue points above the red line). **c-d**, MI values between model predicted M1 spikes and movements (y-axis) versus MI values between recorded mPFC spikes and movements (x-axis), where **c** is for RLPP, and **d** is for SLPP. Different colors of the points represent the M1 neurons from different subjects. The MI values between recorded mPFC spikes and movements is calculated by the average of all mPFC neurons for each subject, since the predictions on each M1 neuron is based on all recorded mPFC neurons. The blue/green line shows the results of linear regression over the data points. The slope of regression line (*k*), correlation coefficient (*r*), and *p*-value are given in each plot. MI values between movements and predictions of RLPP and SLPP both show significant positive correlations with MI values between mPFC spikes and movements, while RLPP has a higher slope, correlation coefficient, and confidence level than SLPP. The dashed diagonal line in red corresponds to the cases when all the motor-related information encoded in mPFC spikes is transferred to the predicted M1 spikes. The MI values for predicted M1 spikes of RLPP and SLPP are both significantly and mostly smaller than the MI values for mPFC (RLPP: *p* = 0.017, 68% of the M1 neurons; SLPP: *p* < 0.001, 97% of the M1 neurons).

### Analyzing the upper bound of behavioral performance for spike prediction models

As aforementioned in Fig. 1, the performance of SLPP should be upper bounded by the correlation between the recordings of the upstream (mPFC) and downstream (M1) regions, while the behavioral information encoded in the recorded upstream signals should determine the upper bound of the RLPP’s performance. We wondered whether these theoretical conclusions could be validated by the real data results, so as to provide insights into the boundary of behavioral performance for spike prediction models.

We first tested the correlation between spike trains from the upstream and downstream (Fig. 8b). When comparing the MI values between spike trains of mPFC and M1 recordings and the MI values between spike trains of mPFC recordings and M1 predictions by SLPP, a strong and significant linear relationship can be found between the compared MI values. Moreover, the majority of the MI values for the SL-predicted spike trains are significantly smaller than the MI values for the real recordings, while the majority of the MI values for the RLPP predictions are significantly higher. These results verify that the performance of SLPP models is upper bounded by the correlation between the recorded mPFC and M1 activities (the x-axis of Fig. 8b), while the RLPP exacts more information from the mPFC recordings.

Then we explored the upper bound for the behavioral performance of RLPP predictions. Fig. **8**c and d reveal how the information encoded in the recorded mPFC signals (x-axis) constrains the behavior-related information captured in M1 predictions (y-axis). An extreme case is when all behavior-related information in recorded mPFC activities is captured by the spike prediction model. Then the MI values between the predicted M1 spikes and movements should be equal to the MI values between the recorded mPFC spikes and movements, shown as the red dashed diagonal lines in Fig. 8c-d. Both RLPP (Fig. 8c) and SLPP (Fig. 8d) show a significant positive correlation below the red line as an upper bound. The difference is, the fitted line of the RLPP has a larger slope, indicating that RLPP can encode more motor-related information than SLPP, closer to the boundary. RLPP also shows a larger correlation coefficient and higher confidence level of the linear relationship. These results suggest that the behavioral performance of the spike prediction models is manipulated by the correlation between recorded upstream spikes and behavior-level signals, while the RLPP can approach the upper limit. The results in Fig. 6k-l also support this claim, where the RLPP performance is robust with respective to decoder settings. Considering that the mPFC activities and target behavioral tasks are the same under different decoders, Fig. 6k-l suggest the direct correlation between the upstream signals and the behavioral performance of RLPP. In other words, if the input for RLPP model is behavior-unrelated, we can expect that the model may fail to generate desired spike patterns.

Collectively, our findings suggest that the RLPP framework can leverage more information conveyed in upstream regions to predict neural spikes in the downstream regions, making it a potentially better choice to generate functional spike trains for behavioral tasks. Our results also point out that the performance of the RLPP framework can still be affected by the quality of the upstream signals.

## Discussion

The hypothesis of this article is that a transregional spike prediction model generates stimulation patterns that can bypass a damaged neural pathway. However, finding a model for the best behavioral performance remains challenging, especially when real activities of the downstream regions are unavailable due to circuit damage. To address this, we introduce here a novel RLPP framework to construct a generative transregional communication model from mPFC spikes to M1 spikes in a behavioral task, a.k.a. a synthetic cortical circuit. The model learns to predict the output spike patterns using behavior-level reward signals via reinforcement learning. We showed that the model predictions on M1 neurons exhibit modulations similar to the real recordings from healthy subjects. Furthermore, the RLPP predictions can be decoded into movements with higher accuracy in the behavioral task than the model trained by real M1 recording via supervised learning. Our analysis suggests that the RLPP model generates more distinguishable spike patterns and leverage more motor-related information from the upstream recordings. This proof-of-concept work highlights the potential of artificially generating neural spike trains from upstream activities as a part of the information pathway among brain regions. We expect that such a framework can be generalized to other neural pathways with proper domain-specific modifications, such as memory, visual prosthesis, *etc*.

The RLPP framework aims not merely to demonstrate the feasibility of spike prediction without recordings of output spikes, but also to surpass constraints of supervised learning-trained models. Our framework offers distinct advantages over SL methods from two perspectives.

First, RLPP allows exploring various output patterns, compensating for model prediction errors and the mismatch between the stimulation and the elicited neural response. The SL-based models aim to reproduce the same neural activities as the real recordings and use such predictions as stimulation patterns. This approach relies on the assumption of one-to-one correspondence between a stimulation on a particular electrode and a spike of an individual neuron^13^. However, evidence suggested that a single microelectrode-delivered pulse would alter the activity of many neurons within a range^26^. This indicates that even the predicted stimulation patterns are the same as recorded spike patterns, the tri ere eural activit o ‘t be the same as desired, and the behavior-level performance is still not guaranteed. Not to mention that the models cannot perfectly predict neural activities when trained by limited recordings. Instead, the RLPP framework is optimized directly toward behavioral performance rather than particular patterns. As shown in our results, RLPP models outperform SLPP models in the behavioral task and are robust to the change of outer environment.

Second, the RL-based transregional model is more bio-plausible in microscopic spike level through behavioral learning and adaptation. The presence of ground truth signals that “teach” euro firing has long been questioned. Instead, brain activities during learning and decision-making are shown to have common computational principles with RL algorithms^27-32^. Our RLPP framework also continuously learns to accomplish the task through trial and error, which makes it open to various new spike patterns and new tasks through reward-guided learning. This means such a framework ca serve as a “silico cortex” participating the information transmission of neural pathways. For example, a larger modulation depth was found in the RLPP predicted spikes, leading to better performance in the behavioral task. This is similar to the neural plasticity observed in subjects learning to control motor BMIs, where the neurons increased the modulation depth or changed the preferred directions^33-38^. External food or water rewards are also effective in changing neural activities and improving the behavioral performance of BMIs^39^, just like our model adapting the spike patterns for more rewards. When changing decoder settings, the RLPP model can generate new patterns distinct from the real recordings, which can be regarded as artificial neurons emerging for new tasks. These observations suggest that the RLPP framework would be more bio-plausible and heuristic for bypassing the damaged neural pathways.

Our method is quite general as each component of the RLPP framework can be replaced and upgraded. The framework consists of a spike prediction model, an RL algorithm for prediction model updating, a behavioral decoder, and an environment to provide rewards. In this article, we used naïve versions of these components, since they already achieved satisfactory results. More advanced tools can be applied to improve the framework’s capability, performance, and efficiency. For instance, using the binless kernel machine^12^ or recurrent neural network^40^ for spike prediction may introduce explainability to the model; the vanilla policy gradient method can be replaced by algorithms like proximal policy optimization^41^, which may lead to faster convergence and better performance. Besides, previous studies showed that internal evaluation of sensory feedback could be estimated from neural activities and applied to RL-based motor BMIs^16, 42-44^. In future work, integrating the internal rewards into the RLPP framework is a potential way toward autonomous model learning.

Particular attention should be paid to the behavioral decoder, which simulates the pathway that generates behavioral responses and further provides rewards to the spike prediction model. The main advantage of using a decoder is to provide a controlled environment in which downstream predictions will always result in timely feedback. For online cases, direct stimulation on the cortical area may not necessarily lead to behavioral changes. Especially in the early training stage, the model predictions can be too inaccurate to evoke behavioral responses. In contrast, the interaction between the decoder and the emulated environment can provide feedback to model predictions at every time instance. Such timely feedback is critical to model training and subject learning, as both the subject and the RLPP model need feedback for adaptation and exploration. Similar paradigms using a controlled environment can be found in muscle stimulation studies^6, 7^. When patients with motor impairments were cued to perform single-joint movements, their muscles were stimulated by electrical pulses predicted from neural signals. At the same time, the patients could see the virtual arm moving on the screen, representing the decoded movements as visual feedback, no matter whether actual arm movements were elicited. A trained decoder can also be helpful for offline training of the spike prediction model, as shown in our study, which may accelerate the learning of subjects.

The decoder also opens a gate of universal design for different subjects as well as manipulating generated spikes patterns. In Fig. 6, we show that the RLPP models can adapt to different decoders and keep the behavioral performance by generating different spike patterns. The major decoder setting used in this study was pre-trained by real M1 spikes and actual movements of the rats. This corresponds to the case that the neural pathway from the target downstream region to the behavior is intact and will be utilized. Here, we did not investigate the dynamics nor the plasticity of the neural pathway. However, literature suggested that subjects could adapt to new decoder settings in motor BMI control in a short time by neural adaptation and reassociation, even with randomly selected parameters^34, 45^. Thus, spike prediction models can actually be trained with a manually designed decoder for different subjects and tuned for better behavioral performance by the co-adaptation between the RLPP model and the neural pathway downward the downstream regions. If necessary, one can manipulate the RL-predictions to desired patterns by carefully designing the decoder. In addition, results of cross-subject decoders provide future directions in transfer learning or brain-to-brain communication.

A recent method, neural co-processor (NCP), adapted the model for predicting stimulation patterns from upstream neural activities with an emulator network to restore lost functions^46^. Our approach is different from this work in the following ways. First, the NCP model is trained by continuous behavior error using supervised learning. However, the error signal requires a ground truth trajectory, which is unavailable after neural pathway damage. Second, the trajectory error needs to be backpropagated through the emulator network to update the NCP model. Thus, the convergence and robustness of the model highly depend on an explicit and accurate emulator network. In contrast, our method directly updates the prediction model from simple behavioral rewards under an RL framework, which is more bio-plausible and does not rely on a specific decoder or emulator. Finally, NCP uses synthetic data generated from simulated neural circuits, while our study is performed on real neural recordings.

The emerging techniques in neural recording and stimulation have revealed the potential of bridging damaged neural pathways for behavioral rehabilitation. Although our study was implemented on mPFC and M1 spike recordings, the RLPP framework provides a general method for generating spike activities from upstream cortical areas and adapting through behavioral feedback. Previous studies have found that Hebbian plasticity could be potentially induced by spike-triggered stimulation^47, 48^. In the long run, the neural stimulation guided by the generated spike trains may also induce cortical rewiring through Hebbian-like plasticity, and eventually lead to long-term rehabilitation beyond temporary function restoration.

## Methods

### Spike Prediction Model

We designed a spike-in, spike-out model for predicting spikes from the upstream brain region to the downstream region (mPFC to M1 in our case). The mPFC spikes are preprocessed into an input ensemble by the Hawkes process^49^. The Hawkes process assumes that past spikes have an exponentially decayed, addictive boosting to current firing probabilities. Given *N*_*x*_ input mPFC neurons and *H* number of past relevant spikes relevant to current time *k*, the input *x*_*k*_ at time *k* is an *H* × *N*_*x*_ matrix, with each element formalized as:

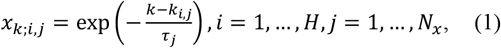

where *k*_*i,j*_ is the timing for the *i*^th^ past spike of the *j*^th^ input neuron; *τ*_*j*_ is the decay parameter for the *j*^th^ neuron. A larger *τ*_*j*_ indicates that past spikes have larger influences on the current firing probability. Compared with the prevailing 0/1 coding for neural spikes, the preprocessed ensemble explicitly describes the information encoded in spike timing and can smooth the input data. In this paper, the decay parameter *τ*_*j*_ is set as 150, and the number of past relevant spikes H is set as 5 for all neurons.

Then, we applied a 2-layer fully connected ANN to capture the nonlinear relationship between the input neural activities and output spikes. The ANN first predicts the firing probabilities of the M1 neurons. Then, the predicted spikes are sampled from the probabilities by an inhomogeneous Bernoulli process. Given *N*_*y*_ as the number of M1 neurons, *λ*_*k,n*_ and *y*_*k,n*_ denote the predicted firing probability and spike of the *n*^th^ output neuron at time *k*, respectively. The *N*_*y*_ length vector ***λ***_*k*_ and ***y***_*k*_ indicate the corresponding values of all output neurons at time *k*. The ANN predicts output spikes as follows:

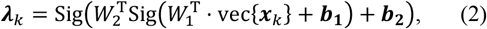

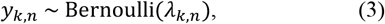

where *W*_1_, *W*_2_, ***b***_**1**_, ***b***_**2**_ are model parameters in compatible size, Sig(·) is the sigmoid function, and vec{·} means vectorization of the matrix. The number of hidden units is set as 64 after exploration. It is worth mentioning that this spike prediction model differs from the conventional classification neural networks. The model predicts spikes as stochastic point process events, not deterministic class labels. Besides, the output layer uses the sigmoid functions on each output node instead of the softmax function over all nodes. This is because each euro ‘s firing is assumed conditionally independent given input firing history.

In order to reduce the output dimension of the spike prediction model, we select the most important M1 neurons related to the task by information theoretical analysis^50^. The M1 neurons are ranked in descending order of the mutual information between the spike trains of M1 neurons and the rat behavior. The top-*N*_*y*_ M1 neurons that have comparable performance in movement decoding as the full M1 neural ensemble are selected as the spike prediction model output, as well as the movement decoder input.

### Movement Decoder and Task Emulator

The movement decoder and task emulator evaluate the predicted M1 spikes and guide the RL optimizer. They serve as a proxy of the natural pathway during the behavioral task: the decoder translates the M1 activities into movements, representing the neuro-muscular pathway from M1 to upper limb muscles; the emulator gives rewards when the decoded movements can accomplish the task, similar to the behavior box providing feedback to the rat. The rewards from the task emulator will be fed to the spike prediction model for RL optimization.

The movement decoder is trained from the recorded M1 spikes and the actual behavior of the rats. In the two-lever discrimination task, we focus on three critical movement states for a successful trial: rest, high press, and low press. We assign the rest movement labels to the time interval of 500 ms before the start cue, and the high/low press movement labels to the duration between the lever press and lever releasing of the rats. A 2-layer fully connected ANN is trained to decode the M1 spikes into these three movements. The input ***d***_*k*_ is an (*N*_*y*_ × *H*_*d*_) length vector defined as vec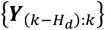, where ***Y***_*k*_ represents the recorded spikes of M1 neurons at time *k*, and *H*_*d*_ is the history length of M1 spikes considered by the decoder, chosen to be 60, which covers 600 ms history of M1 spikes to include enough past information. The ANN outputs a 3-element vector corresponding to the three movements, and the one with the highest output value is the decoded movement. The decoder can be formalized as:

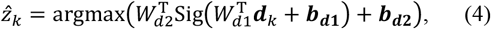

where 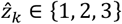 is the index of the three movements. The number of hidden units *N*_*d*_ is explored separately for each rat. The decoder parameters, *W*_*d*1_, *W*_*d*1_, ***b***_***d*1**_, ***b***_***d*2**_ are optimized by the cross-entropy using the scaled conjugate gradient method^51^. The data samples are randomly divided into train, validation, and test sets by a ratio of 70%, 15%, and 15%, respectively. The model training is stopped when the performance on the validation set gets worse to avoid overfitting. The weights of the ANN are randomly initialized multiple times to find the optimal decoder. The weights of the movement decoder are fixed when training the spike predictions model.

The task emulator is designed to evaluate the interaction of decoded movements with the environment. Given the predicted M1 spikes, the emulator would return a reward *r*_*o*;*k*_ = 1 if the movement decoded from the predicted M1 spikes fits the task requirement, and *r*_*o*;*k*_ = 0 if the movement fails to do so. The subscript *o* emphasizes that *r*_*o*;*k*_ is an outer reward from the environment. Unlike the behavior box, which only gives feedback to the rat at the end of the trial, the emulator provides instantaneous feedback every 10 ms, as the movement is continuously decoded from the M1 spikes at each time bin.

### Reinforcement-based Model Optimization

We used the policy gradient algorithm to optimize the spike prediction model with the behavioral reward. According to the policy gradient theorem^52^, a pseudo gradient of model parameters can be estimated by a Monte Carlo method. The model parameters ***θ*** = vec{*W*_1_, *W*_2_, ***b***_**1**_, ***b***_**2**_} is then updated by the mini-batch gradient descent with a maximum iteration number of 5000. The model weights with the best success rate in the behavioral task out of all the iterations would be used for testing. In each training iteration, the parameters are updated as below:

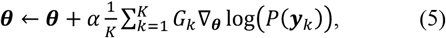

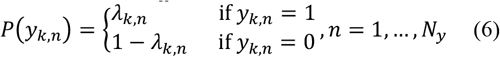

where *α* is the learning rate, *K* is the number of time bins in one mini batch, ∇_***θ***_ means taking the derivative with respect to ***θ***, *G*_*k*_ is the cumulative discounted return at each time instance within one trial, and the *P*(***y***_*k*_) is a *N*_*y*_ length vector representing the firing probability of ***y***_*k*_. During training, the learning rate *α* linearly decays from 1.2 to 0.5 as the number of iterations increases to avoid overfitting. The model performances are assessed with the five-fold cross-validation. Data is randomly divided and shuffled by trial during training. Each mini batch contains data samples from 20 successful trials. The model is re-initialized 32 times, and the one with the best performance is used for further analysis.

The cumulative discounted return *G*_*k*_ is a behavior-level evaluation of the predicted M1 spike patterns. It considers that current spikes have a cumulative, time-decayed contribution to future rewards. We normalize *G*_*k*_ by a z-score function to emphasize the spike patterns that lead to better behavioral performance in a mini-batch and penalize the rest. Formally, *G*_*k*_ is expressed as:

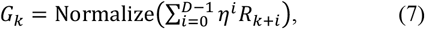

where *D* is the delay time length and *η* ∈ [0, 1] is the discount factor. A large *η* indicates that the current spike predictions are assumed to have a long-term effect on future behavior. Here, we set *D* and *η* to be 100 and 0.98 after parameter exploration, which assumes that all future rewards received within 1 second are contributed by current spikes on a decayed basis. The *R*_*k*+*i*_ is the immediate reward of the behavioral output at time *k* + *i*, which should be appropriately designed to achieve desired behavioral performance.

### Reward design and prior knowledge for RL training

The trade-off between exploration and exploitation is critical for the efficiency and convergence of RL training. In practice, simply using the vanilla policy gradient with outer rewards from the task emulator can lead to poor performance, as the models often converge prematurely, hindering their ability to fully explore potential spike patterns. Here, we introduce two operations to avoid such scenarios: inner reward design and spike generation with prior knowledge.

In reward design, we enrich the reward function *R*_*k*_ with an inner reward term for behavior-level exploration. Different from the outer reward *r*_*o*;*k*_, which reflects the success of the task, we design the inner reward *r*_*i*;*k*_ to encourage the exploration on rarely encountered movements and keep the “inner curiosity” of the model. For instance, when the low lever press dominates the decoding results in a given training iteration, we will set a larger inner reward for the high lever press and rest movements. A factor *ε* is used to balance the outer and inner reward:

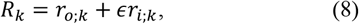

where *∈* linearly decayed from 1 to 0 during training to prevent the model from being biased by the inner reward.

Generating spikes with prior knowledge is used for the exploration of single neurons firing. We notice that the RLPP model tends to be trapped in the local optimum where the predicted firing probabilities are constantly close to 0 or 1, causing gradient vanishing and poor performance. In contrast, our empirical knowledge suggests that the models can better adapt to different inputs when the predicted firing probability has a mean roughly between 0-0.5 and a relatively high variance. Thus, we force the model to explore more within the desired firing range, especially in the early training stage. Specifically, the predicted spikes are generated from a Bayesian correction of model predictions *λ*_*k,n*_, written as 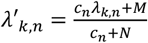. The latter part 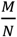 controls the prior estimation of the firing probability, which is set as 0.13, close to the average firing probabilities of real M1 recordings. The confidence level of current model predictions, *c*_*n*_, increases as the number of training iterations and the variance of the predicted firing probabilities increase.

### Supervised-based Model Optimization

The spike prediction models are also trained by supervised learning to compare with the RLPP framework. The SL training has the same procedure as the RL training, but uses the mini-batch gradient descent with five-fold cross-validation. The only difference is the loss function. In supervised-based model optimization, the model parameters are optimized by maximizing the point process log-likelihood function between the model predictions and the recordings of M1 spikes. The iteration is written as below:

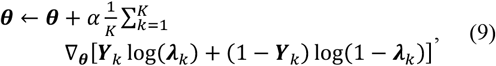

where *α* starts from 1.1 and linearly decays to 0.1 as the number of iterations increases.

## Notes

### Competing Interest Statement

The authors have declared no competing interest.

